# Consumption of clinically relevant cannabidiol doses during pregnancy alters postnatal behavior in offspring

**DOI:** 10.1101/2025.04.30.651495

**Authors:** Luis E Gomez Wulschner, Victoria N Chang, Won Chan Oh, Emily Anne Bates

**Author notes:** Co-corresponding authors **COORESPONDANCE TO:** Emily Anne Bates, PhD, University of Colorado Anschutz Medical Campus, 12800 E 19^th^ Avenue, RC1 North, Box 8313, Aurora, CO 80045, Won Chan Oh, PhD, University of Colorado Anschutz Medical Campus, 12800 E 19^th^ Avenue, RC1 North, Box 8303, Aurora, CO 80045. Authors contributed equally (Luis E Gomez Wulschner, Victoria N Chang).

## Abstract

Pregnant people use cannabidiol (CBD), a non-psychoactive cannabinoid of cannabis, due to its perceived safety and to treat side-effects such as nausea, insomnia, pain, and anxiety. However, CBD crosses the placenta and accumulates in the fetal brain, where it can activate or repress the function of several molecular targets that are important for brain development. While consumption of high doses of CBD during pregnancy have been shown to disrupt offspring neurodevelopment and postnatal behavior, lower, more clinically relevant doses have not been assessed. Here, we show that oral consumption of 10 mg/kg/day CBD during pregnancy increases thermal pain sensitivity in exposed male mouse offspring. Additionally, we find that the same dose impairs cognition and reduces excitability of the prefrontal cortex in exposed female mouse offspring. These data show that lower doses of CBD consumption during pregnancy can impair fetal brain development and postnatal behavior.

## INTRODUCTION

Pregnancy is often accompanied by many unpleasant and debilitating side effects such as nausea, pain, insomnia, and anxiety^1^. To counter these effects, many pregnant people self-medicate with cannabis due to its antiemetic and anti-anxiety properties^1^. Cannabis is composed of two primary cannabinoids, tetrahydrocannabinol (THC) and cannabidiol (CBD). THC is typically known as the psychoactive component in cannabis with legal access being restricted in parts of the United States and some other countries^2^. CBD, however, is viewed as non-psychoactive and is federally legal in many parts of the world^2^. Based on a study of self-reported cannabis use among pregnant women from 2009 to 2017, the adjusted prevalence of use increased from 6.80% to 12.50%^3^. This percentage only appears to be increasing with the self-reported prevalence of cannabis consumption among pregnant women in the United States now up to 26%^4^. Additionally, umbilical cord blood samples reflect a similar prevalence of THC consumption, showing that 22.4% of pregnant women had detectable THC metabolites present^5^. These assessments, however, do not evaluate the consumption of CBD-only products. Given CBD’s touted therapeutic effects and lack of associated social stigma, a segment of the population will readily consume federally legal CBD, even if they do not consume whole cannabis or THC. For similar reasons, many believe it to be safer to use over other drugs during pregnancy. In a recent study assessing CBD use during pregnancy, the prevalence of CBD-only use in pregnant women reached 20% which was significantly higher when compared to non-pregnant CBD using women^6^. Despite its perceived safeness, very little is known about the consequences of prenatal CBD exposure on fetal development.

CBD can cross the placenta, accumulating in the developing fetal tissue and acting upon different targets relevant to fetal brain development ^7,8^. Furthermore, cannabis consumption during pregnancy is associated with neurodevelopmental disorders during adolescence, suggesting that exposure to CBD alone could be partly responsible for some of these behavioral effects^9,10^. Many of these developmental behaviors are mediated by the pre-frontal cortex (PFC), a region of the brain shown to be dysregulated following exposure to cannabis and CBD ^11,12^. The PFC expresses many receptors that CBD is capable of affecting, many of which have been implicated in neuronal developmental processes such as neurogenesis and synapse development ^13–16^. Therefore, the high expression of CBD targets in the fetal brain suggests that exposure to CBD during pregnancy could impact fetal brain development and postnatal behaviors We have previously shown that administration of high dose (50mg/kg/day) CBD to C57BL6 female dams results in sex-specific effects, with increased thermal sensitivity in male CBD-exposed offspring and cognitive deficits in female CBD-exposed offspring. Furthermore, the same dose resulted in reduced excitability of pyramidal neurons in the PFC of the female CBD-exposed offspring^17^. However, whether lower doses of CBD disrupt fetal brain development and postnatal behavior remained an open question. Here we use a lower, more clinically relevant dose (10mg/kg/day), to assess whether these changes in behavior and neuronal activity are maintained. Due to the sex-specific effects seen in our previous research, in this study we evaluated male CBD-exposed offspring for thermal sensitivity and tested female CBD-exposed offspring for PFC associated tasks and neuronal mechanisms. We found that gestational exposure to 10 mg/kg/day CBD increased thermal sensitivity in male offspring and caused PFC-associated cognitive deficits and changes in excitability in female offspring. Together, this suggests that chronic clinical doses of CBD during the prenatal period could result in adverse effects on fetal development.

## RESULTS

### 10 mg/kg/day CBD exposure does not affect gestational length, litter size, or survival

CBD is most consumed orally or through topical application and so to model human consumption, we administered CBD or vehicle by oral gavage^6^. We administered 10mg/kg of CBD dissolved in sunflower oil or sunflower oil alone (control) daily via oral gavage to C57BL6J female pregnant dams from E5 to birth. To determine how this dose of CBD administered during pregnancy affects maternal factors, we quantified the size of litters, gestational weight gain, and gestational length of both the control and CBD dosed dams and litters (Fig 1A). Compared to the sunflower oil alone condition, CBD consumption during pregnancy did not significantly change any of these maternal or litter factors (Fig. 1B).

**FIGURE 1:**
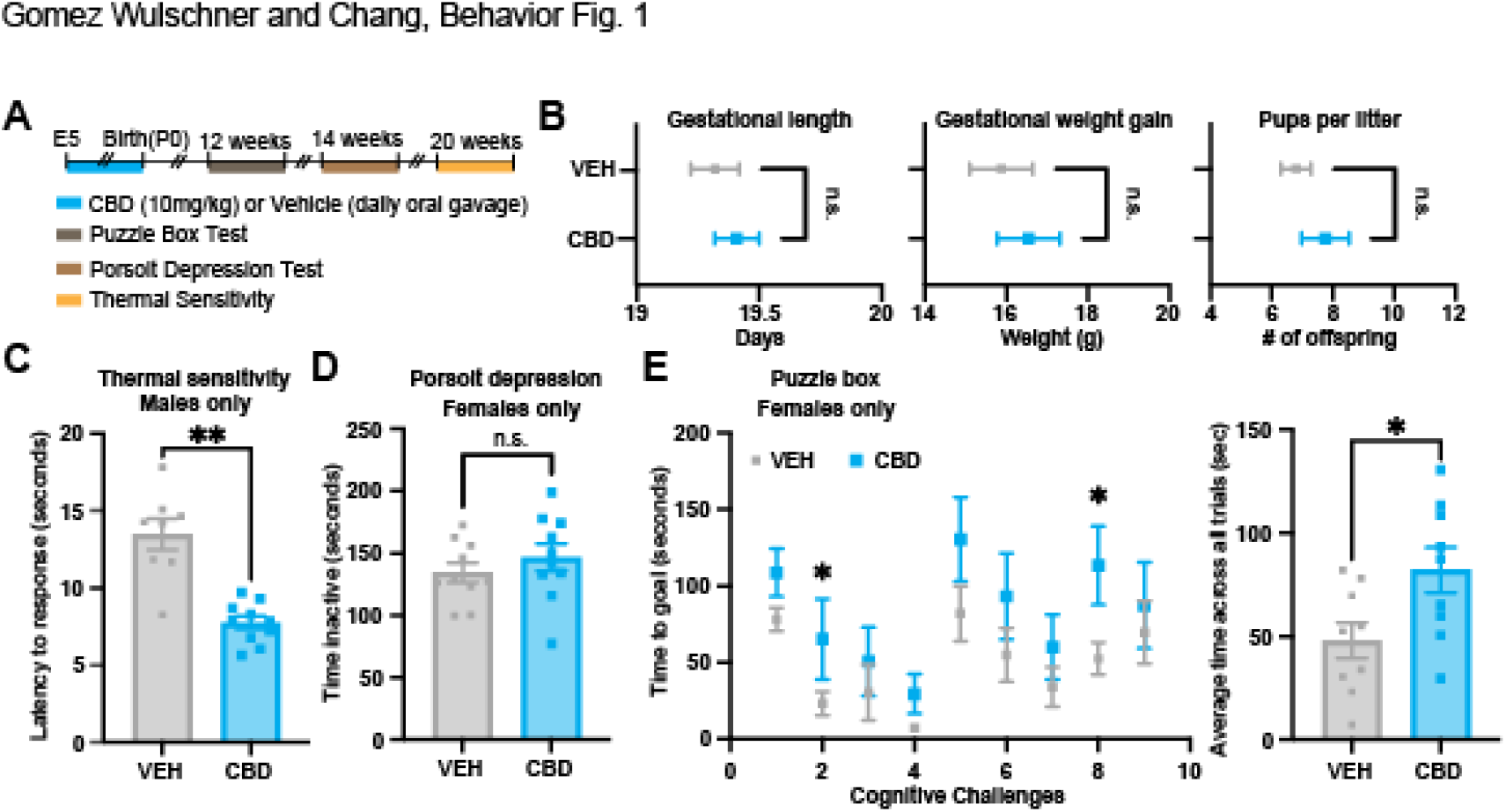
Fetal CBD exposure alters postnatal behavior in a sex specific manner. **A**, Experimental timeline for fetal CBD and vehicle treatment (daily oral gavage) and postnatal behavior paradigms. **B**, Dams’ gestational length (p=0.54, t-test), gestational weight gain (p=0.57, t-test), and litter size (p=0.296, t-test) were unaffected by CBD (blue squares) compared to vehicle (gray circles) treatment (gestational length and weight, n=18 vehicle and 11 CBD; litter size, n=10 vehicle and 8 CBD). **C**, The Hargreaves test displayed increased thermal sensitivity in fetal CBD exposed male offspring (vehicle: n=8 male mice; CBD: n=10 male mice; p<0.0001, t-test). **D**, The Porsolt forced swim depression test showed no difference in vehicle and CBD treated female offspring (vehicle: n=10 female mice; CBD: n=10 female mice; p=0.38, t-test). **E**, Performance in puzzle box test was impacted in CBD exposed female offspring (vehicle: n=9 female mice; CBD: n=9 female mice; p= 0.02, t-test). ^*^P<0.05, ^**^P <0.01; error bars represent SEM. n.s. not significant.

### 10mg/kg/day CBD prenatal exposure increases thermal pain sensitivity in male offspring

CBD is a promiscuous drug, binding to many different receptors important for proper neuronal development including the heat-activated transient potential vanilloid receptor one (TRPV1) calcium channel^13^. In our previous study, we showed that 50mg/kg/day of CBD during prenatal development led to heightened responses of male offspring to the Hargreaves test^17^. The Hargreaves test measures latency to respond to thermal stimuli and thermal pain sensitivity^18^. Because the higher dosing of CBD increased thermal sensitivity in males, we tested whether prenatal 10mg/kg CBD daily exposure was sufficient to elicit the same effect. Indeed, we found that latency to respond to the thermal stimuli was significantly faster in males exposed to CBD during pregnancy (Fig. 1C). This suggests that lower, more clinically relevant doses of CBD during pregnancy are sufficient to increase thermal sensitivity in males.

### 10mg/kg/day CBD prenatal exposure does not result in depression-like behaviors in female offspring

There is previous evidence to suggest that prenatal exposure to cannabis results in higher rates of anxiety, depression, and ADHD during adolescence^9,10^. Despite not observing any changes in anxiety-like behaviors in our previous study, we did not investigate whether there were any changes in depressive-like behaviors following prenatal CBD exposure. Therefore, we assessed whether female offspring exposed to 10mg/kg/day CBD during pregnancy had increased depression-like behaviors in the Porsolt forced swim depression test^19^. We found that CBD exposed females did not have a significantly longer time spent inactive compared to controls (Fig. 1D), suggesting that prenatal CBD exposure does not result in increased depressive-like behaviors.

### 10mg/kg/day CBD prenatal exposure decreases problem-solving abilities female offspring

Another possible long-term consequence of prenatal CBD exposure is its effects on childhood and adolescent cognitive abilities such as problem solving^9^. Indeed, we found that female offspring exposed to higher dose CBD (50mg/kg/day) exhibits impaired problem-solving behavior^17^. To evaluate whether the 10mg/kg/day CBD exposure has a similar impact on problem-solving abilities, CBD-exposed female offspring or control mice underwent the puzzle box test. During this test, mice are in a lit box environment and are presented with progressively harder problem-solving tasks in order to reach a dark box goal environment^20^. Each mouse completes nine trials, with novel progressive challenges at trial two, five, and eight. Mice that are capable of problem-solving are quicker to reach the dark environment upon secondary exposure to the task^20^. Female offspring exposed to 10 mg/kg CBD throughout gestation took significantly more time to reach the goal area in trial two and eight compared to control mice (Fig. 1E). CBD-exposed female offspring took more time to reach the goal area across trials, suggesting that this lower dose of CBD exposure impairs problem-solving behaviors in female mice.

### 10mg/kg/day CBD prenatal exposure decreases excitability of layer 2/3 pyramidal neurons in the PFC of female offspring

The PFC is a brain region critical for higher-order cognitive behaviors^21,22^. Disruption of normal PFC function during early development has been implicated in several neurodevelopmental disorders^21–23^. Due to the effect of CBD on problem-solving behaviors, we investigated the neuronal mechanisms underlying this impairment. We previously showed a higher dose of CBD (50mg/kg/day) decreased excitability of PFC layer 2/3 pyramidal neurons only in female offspring^17^. To determine whether this effect remained following 10mg/kg prenatal CBD daily exposure, we electrophysiologically measured the intrinsic membrane properties of layer 2/3 pyramidal neurons in acute PFC slices (Fig. 2A). Indeed, 10mg/kg/day prenatal CBD significantly decreased excitability and increased the minimum current required to trigger an action potential compared to female controls without affecting the resting membrane potentials (Fig 2. B-E). This suggests that 10mg/kg/day CBD daily exposure during pregnancy is sufficient to induce changes in PFC neuronal excitability in females.

**FIGURE 2:**
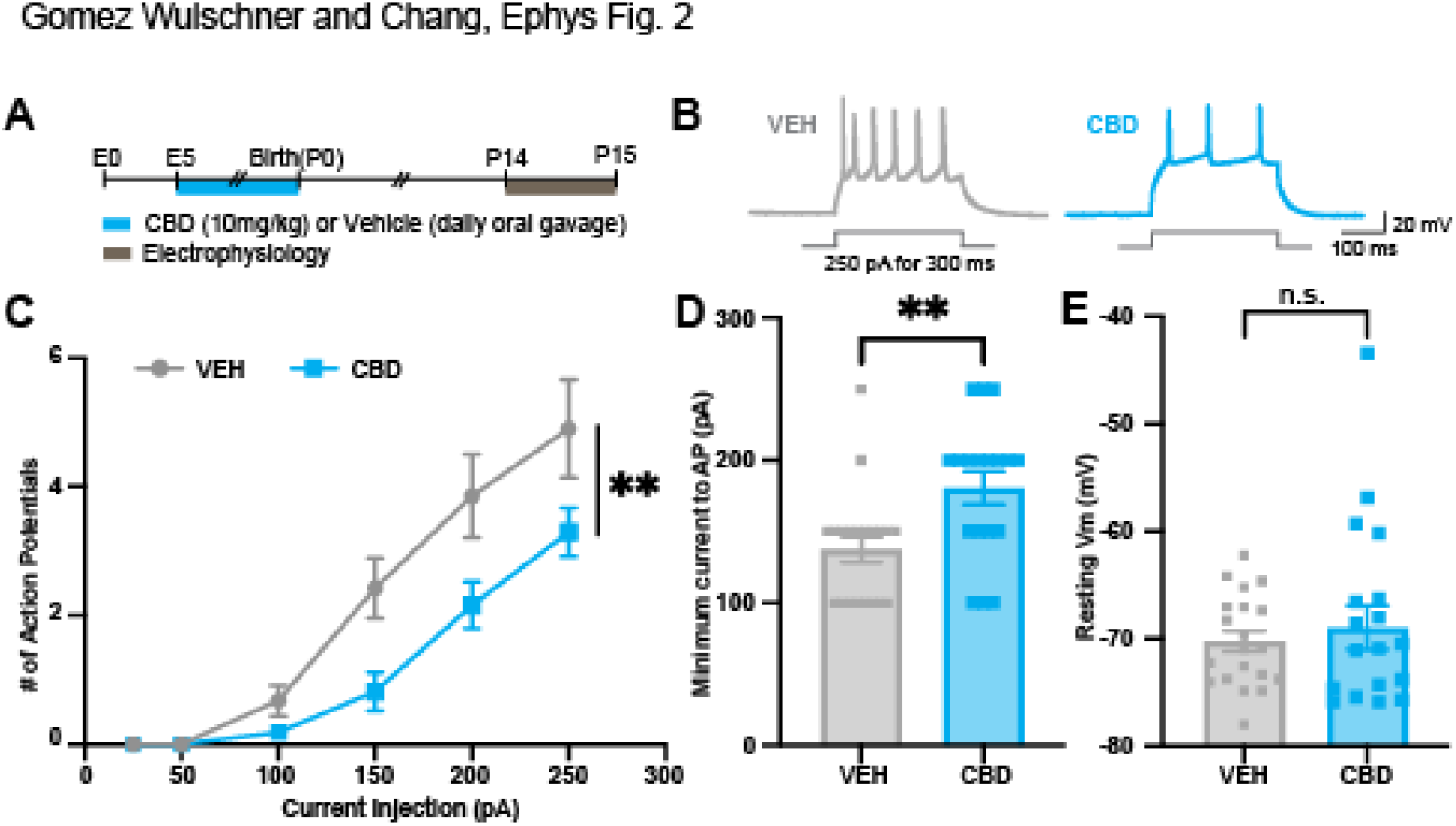
Fetal CBD exposure decreases excitability of layer 2/3 pyramidal neurons in female offspring. **A**, Experimental timeline of fetal CBD exposure (daily oral gavage) and postnatal electrophysiology. **B**, Representative traces of an action potential train elicited by 250 pA current injection for 300 ms in vehicle (gray) and CBD (blue) exposed female offspring. **C**, Fetal CBD exposure significantly reduced intrinsic excitability of PFC layer 2/3 pyramidal neurons compared to vehicle exposed controls (vehicle: n=20 cells, 3 mice; CBD: n=19 cells, 3 mice; treatment effect: p <0.0001, 2-way ANOVA). **D**, The minimum current required to elicit action potential spikes was increased in CBD exposed female offspring (vehicle: n=20 cells, 3 mice; CBD: n=18 cells, 3 mice; p=0.0049, Mann-Whitney test). **E**, Resting membrane potential was unaffected by CBD (vehicle: n=20 cells, 3 mice; CBD, n=19 cells, 3 mice; p=0.7919, Mann-Whitney test). ^*^P<0.05, ^**^P <0.01; error bars represent SEM. n.s. not significant.

## DISCUSSION

CBD use during pregnancy has become of increasing concern in large part due to a rise in accessibility as well as its perceived safety^6^. Here, our data demonstrate that exposure to clinically relevant levels of CBD (10mg/kg/day) during pregnancy results in increased thermal sensitivity in male offspring and decreased problem-solving behaviors and excitability of PFC in female offspring in mice.

CBD binds to many receptors including the heat sensitive TRPV1 receptor. Increased thermal sensitivity of males indicates that fetal CBD exposure results in a heightened sensitivity of these receptors even at this lower dose. Therefore, even low levels of CBD during fetal development are capable of impacting TRPV1 thermal pain circuits potentially leading to negative long-term behavioral effects^24^.

The PFC is a critical region of the brain not only for higher-order cognitive behaviors, but its proper function is crucial for normal brain development^21,22^. Clinical levels of CBD affect proper PFC development resulting in decreased problem-solving behaviors in female offspring. CBD is also an agonist for the serotonin receptor 5-HT1A (5-HT1ARs). One potential neuronal mechanism through which CBD could be eliciting this effect is through binding to inhibitory 5-HT1ARs. Activation of 5-HT1ARs has been shown to decrease neuronal activity and are not only activated by CBD but are also highly expressed in the PFC during early development^25–27^. We observed that 10mg/kg/day of CBD during fetal development leads to decreased excitability of layer 2/3 pyramidal neurons in the PFC. Thus, exposure to CBD during pregnancy could potentially aberrantly activate 5-HT1ARs, thereby leading to decreased PFC excitability and deceased problem-solving abilities. Overall, the underlying mechanisms for the sex-specific effects seen here are yet to be uncovered and future studies will aim to investigate how fetal CBD exposure leads to neurodevelopmental consequences.

Altogether, these results show that even at lower, more clinically relevant levels, CBD exposure during pregnancy results in sex-specific, adverse neurodevelopmental effects in mice.

## METHODS

### Mice

Eight-week-old female C57Bl6 (Strain #000664) mice from Jackson laboratories (Maine) were crossed with male C57Bl6 mice from the same source. Upon observation of a vaginal plug (embryonic day 0.5), mice were individually housed for the duration of pregnancy. Dams were weighed and administered 10 mg/kg/day CBD dissolved in sunflower oil or sunflower alone (vehicle) by oral gavage starting on embryonic day 5 through birth. Pups were housed with their mother until they were weaned and separated by sex on postnatal day 21. Mice were fed a standard chow diet. The University of Colorado, Office of Laboratory Animal Research (OLAR) oversees an AAALAC accredited animal facility that meets NIH standards as outlined in “Guide for the Care and Use of Laboratory Animals”. The institution also accepts as mandatory PHS “Policy on Humane Care and Use of Laboratory Animals by Awardee; Institutions and NIH “Principles for the Utilization and Care of Vertebrate Animals Used in Testing, Research, and Training”. An approved Assurance of Compliance is on file for Protection from Research Risks. Experiments were approved by the University of Colorado Anschutz Medical Campus Institutional Animal Care and Use Committee (protocols #139 and 721).

### Preparation of acute prefrontal cortex (PFC) slices

Acute coronal PFC slices were obtained from P15 C57BL/6 female wild-type mice prenatally exposed to either CBD (10 mg/kg/day) or vehicle. Mice were anesthetized with isoflurane and euthanized by decapitation. Immediately after decapitation, the brain was extracted and placed in icy cutting solution containing (in mM) 215 sucrose, 20 glucose, 26 NaHCO3, 4 MgCl2, 4 MgSO4, 1.6 NaH2PO4, 1 CaCl2, and 2.5 KCl. Using a Leica VT1000S vibratome, the PFC was sectioned into 300μm thick slices. PFC slices were incubated at 32°C for 30 minutes in 50% cutting solution and 50% artificial cerebrospinal fluid (ACSF) composed (in mM) of 127 NaCl, 25 NaHCO3, 25 D-glucose, 2.5 KCl, 1.25 NaH2PO4, 2 CaCl2, and 1 MgCl2. After 30 minutes, this solution was replaced with ACSF at room temperature. For all electrophysiology experiments, the slices were placed in a recording chamber and bathed in recirculating ACSF aerated with 95% O2 / 5% CO2 at 30°C and allowed to equilibrate for at least 30 minutes prior to the start of experiments.

### Electrophysiology

PFC layer 2/3 pyramidal neurons were identified by morphology and distance from layer 1 and patched in the whole-cell (8-10 MΩ electrode resistance; 20-40 MΩ series resistance) current clamp configuration (MultiClamp 700B, Molecular Devices). Using a potassium-based internal solution (in mM: 136 K-gluconate, 10 HEPES, 17.5 KCl, 9 NaCl, 1 MgCl2, 4 Na2-ATP, 0.4 Na-GTP; ∼300 mOsm, and ∼pH 7.26), spiking properties were examined at 30°C in recirculating ACSF with 2mM CaCl2 and 1mM MgCl2. Excitability was measured by injection of depolarizing current steps (25 - 250pA, 300ms). Resting membrane potential was recorded prior to the first depolarizing current step. The minimum current required to elicit an action potential was defined as the smallest current step triggering at least one spike (PMID: 37433966).

### Behavior tests

#### Puzzle Box Test

Female mice underwent the puzzle box test at 12 weeks of age. The puzzle box is performed in a box divided by barrier into two compartments: a brightly-lit start zone and a smaller covered goal zone. Mice are motivated by their aversion to the bright light to reach the goal zone. Mice undergo nine trials (T1 toT9) over 3 consecutive days in which they are challenged with increasingly difficult obstructions. This sequence assesses problem-solving abilities (T2, T5, and T8), learning/short-term memory (T3, T6, and T9), and repetition on the next day measures long-term memory (T4 and T7).

#### Porsolt Forced Swim Test

Mice underwent the Porsolt forced swim test at 14 weeks of age. The forced swim test is the most widely accepted model of depression-like behavior in mice. It is performed only once per animal. The characteristic behavior of the test, termed immobility, develops when a rodent is placed in a tank of water for an extended time and “makes only those movements necessary to keep its head above water”^28^. Forced swim sessions are conducted by placing mice individually in large glass cylinders (45 x 20 cm) containing 24-25 C° water approximately 20 cm deep. The water should be at a height such that the mouse cannot escape or touch the bottom of the beaker. The mouse is placed in the cylinder for 6 min. Latency to float and amount of time spent struggling are measured. Trials are video recorded and later scored by a blind observer. During the entire duration of the task, the experimenter is present and watching the mice. If there is any indication that the animal is in danger of drowning, it is immediately removed from the cylinder and excluded from the study. At the end of the swim session the mice are towel dried and returned to their home cage. The water in the test arena is changed between each subject.

### Hargreaves Thermal Sensitivity Test

The Hargreaves thermal sensitivity test measures latency to response in seconds to a calibrated heat source (light) that is directed at the center of each individual mouse hind paw. Removal of the paw or flinching is recorded as a response. Measurements were taken three times per paw for each mouse and averaged together.

#### Statistics and Reproducibility

All experiments include a minimum of 3 mice per group in the behavioral paradigms and acute prefrontal cortex preparations. Statistics were calculated across mice or cells and reported on figure legends. All error bars represent standard error of the mean and alpha value was set at 0.05 for all experiments. Datasets were compared using an unpaired two-tailed t test, Mann-Whitney test, or two-way ANOVA depending on normality, equal variance, and experimental design. All datasets include outliers. Single and double asterisks indicate P < 0.05 and P < 0.01, respectively. n.s., not significant.

## ACKNOWLEDGMENTS

We thank the Animal Behavior Core for conducting depression and puzzle box experiments. We would like to thank the Institute of Cannabis Research, Colorado for funding to Emily Bates for this project. We would like to thank the Ludeman Center for Women’s Health Research for funding Won Chan Oh and Emily Bates for this project. This work was also supported by NIH R01MH124778, R21MH137409, and R21NS133681 (W.C.O.). We are grateful for this funding, without which this work could not have been completed.

